# Normative models reveal distinct cortical abnormalities to dimensions of psychopathologies in preadolescents

**DOI:** 10.1101/2024.11.29.626043

**Authors:** Qingkun Deng, Elizabeth Levitis, Rick A Adams, Andre Altmann

## Abstract

**Background:** Evidence suggests a non-specific mapping between psychiatric disorders and underlying neurobiological substrates. A dimensional psychopathology framework may prove useful for organizing observed neurobiological alterations along broad psychopathological dimensions.

**Methods:** We applied latent class analysis, with an additional constraint on classification uncertainty, to identify clinical cohorts of symptomatic homogeneity to represent the high-risk end of specific psychopathological dimensions (i.e., internalizing/externalizing, *p-factor*), using baseline data (*N* = 11860) from the Adolescent Brain and Cognitive Development (ABCD) Study. These cohorts were compared against neurotypical individuals in deviations from the normality of cortical development, quantified using autoencoder-based normative models, to reveal cortical abnormalities.

**Results:** We identified cortical thickness related to psychopathologies in the ABCD data, particularly to externalizing syndromes, and revealed distinct structural abnormalities to broad psychopathological dimensions.

**Conclusion:** This study highlights the value of person-centered analytic techniques, combined with normative modeling, to complement traditional associational methodologies in revealing neurobiological correlates of dimensional psychopathologies.

## Introduction

To date, most psychiatric neuroimaging studies have been conducted within a case-control study framework, using psychiatric diagnoses to define clinical groups and assuming that neurobiological alterations are uniformly shared within disorders and are distinct between them. This assumption, however, has been disputed by large-scale meta-analyses which have found few diagnostically specific brain morphological changes (1). Instead, common structural changes, such as grey matter reductions in the dorsal anterior cingulate and insula (1), implicated in functional alterations within and between major neurocognitive networks, are found to be shared across psychiatric disorders (2,3). Cumulative evidence, therefore, suggests a non-specific mapping between diagnostic categories and underlying neurobiological abnormalities.

Such a non-specific mapping may be expected given the high rates of psychiatric co-morbidity (e.g., about 40% of adolescents impacted by mental illness have multiple diagnoses (4)), which suggests unitary psychopathological conditions or constructs being divided into arbitrary categories (5). Indeed, a large number of symptom-level factor analytic studies have empirically identified transdiagnostic psychopathological dimensions that explain the systematic co-occurrence of symptoms across diagnostic categories (6–8). Evidence from these studies led to the development of dimensional nosology of psychopathology, such as the Hierarchical Taxonomy of Psychopathology (HiTOP) framework (5), conceptualizing psychopathological conditions as continua with normal to clinical variations, instead of categorical entities.

Two psychopathological dimensions — internalizing and externalizing — are supported by substantial evidence in both adults and adolescents (9). The internalizing dimension reflects an individual’s distress expressed inwardly, involving syndromes typically seen in mood disorders such as major depressive disorder and generalized anxiety disorder. The externalizing dimension, on the other hand, reflects overt behavior rendering an individual in social conflict with others, incorporating syndromes seen in, for instance, oppositional defiant disorder and conduct disorder. The two dimensions are moderately and positively correlated with each other (8), which can be accounted for by a higher-order factor of general psychopathology (i.e., *p-factor*) (10), representing a shared susceptibility for developing any mental disorders (11).

A dimensional nosology, as an alternative to the traditional taxonomy, may prove useful for unifying observed neurobiological alterations, non-specifically associated with different disorders, along broad psychopathological dimensions (8). Methodologically, dimensional modeling of psychopathologies as continuous variables (e.g., factor scores derived from factor analysis) allows an association analysis (e.g., correlative, regression-based) of the relationships between brain-derived features and psychopathologies. However, while some studies adopting a dimensional approach reported dissociable neurobiological correlates to specific psychopathological dimensions (e.g., fear, internalizing, externalizing) (12–16), others found very few neurobiological correlates unique to distinct psychopathological dimensions and revealed widespread, non-specific, alterations in brain structures associated with higher *p-factor* scores (17–20).

The methodological emphasis on continuous associations has led much of the prior research to identify linear, often univariate, brain-behavior relationships, capturing broad variations across both clinical and non-clinical ranges in behavioral variables. While a simple linear assumption facilitates a straightforward interpretation, such methods disregard the potentially non-linear, non-monotonic, associations that may underlie brain-behavior relationships and relationships between brain-derived features. For instance, if neurobehavioral relations manifest differently between normative and clinically relevant ranges of behavior, neurobiological correlates identified in the previous association studies, involving large non-clinical populations, could be non-characteristic of psychopathologies and therefore non-specific to broad psychopathological dimensions.

Here, we evaluate whether Latent Class Analysis (LCA) (21), a person-centered analytic technique that identifies latent groups of individuals with phenotypical similarities, may provide a valuable tool for the study of dimensional psychopathologies and their neurological correlates. Specifically, we add an additional constraint on classification uncertainty to LCA to isolate cohorts of relatively homogeneous behavior syndromes in a large cohort of preadolescents from the Adolescent Brain and Cognitive Development (ABCD) Study. We hypothesize that these cohorts, with reasonably high symptomatic homogeneity and clinical relevance, may serve as relatively precise targets to represent the high-risk end of psychopathological dimensions for which specific neurobiological abnormalities can be ascribed.

We focus on the structures of the cerebral cortex and study their abnormalities via *normative modeling (22)*. A normative model maps the individual variations of cortical structures in normative, neurotypical individuals with no behavioral syndromes. Cortical abnormalities of individuals with psychopathologies are, therefore, understood in relation to these variations, quantified as deviations from an expected pattern of normative neurodevelopment. Instead of conventional normative modeling methods (e.g., Gaussian process regression), which establish normality for individual brain features in isolation, we apply autoencoder-based normative models (23) to account for the potentially non-linear spatial interactions between cortical regions, together with squared, non-directional measurement of deviations, without assuming a uni-directional, monotonic manifestation of cortical abnormalities in individuals with similar psychopathologies. We derive cortical abnormalities of specific psychopathological dimensions via group comparisons between clinical cohorts against neurotypical individuals in terms of their quantified deviations from normality. Our approach may provide higher sensitivity for identifying neurobiological abnormalities, compared to linear association analysis and traditional case-control analysis, for its assumption-free modeling of the data and group comparisons in deviations instead of feature values themselves.

## Materials and method

### Study sample

The ABCD study is a longitudinal study consisting of a demographically diverse sample of over 11,000 children, aged 9-10 at baseline assessment, probabilistically sampled from schools around 21 research sites across the United States (US) (https://abcdstudy.org/). The recruitment process was detailed in Garavan et al. (24). Ethical review and protocol approval were secured from the Institutional Review Board (IRB) at the University of California, San Diego, and from local IRB approval (25). Parents or guardians gave written informed consent after receiving a full explanation of the procedures, and children provided assent before participating in the study. We used the baseline data from the ABCD Study’s Data Release 5.1 (*N* = 11868). We considered all subjects with complete information for the Child Behavior Check List (CBCL) measures (26) (*N* = 11860) for performing LCA, described below. Sample characteristics are presented in the Supplement (Table S3, Figure S2). For neuroimaging analysis, we excluded intersex-male subjects (*n*= 3), subjects whose scans failed to pass quality control by the recommended T1-weighted imaging data inclusion criteria or considered by neuroradiologists for clinical referral (*n* = 916), and subjects with incomplete data for CBCL syndrome scales (*n* = 6) and psychiatric diagnoses (instruments described below, *n* = 113). A final neuroimaging analysis sample of *N* = 9021 unrelated subjects (randomly selecting one subject from each family) was available for normative modeling analysis, described below.

### MRI image acquisition and processing

Full details of the MRI acquisition protocols and processing procedures can be found elsewhere (27,28). Briefly, MRI data were collected using 3T scanners, specifically the Siemens Prisma, GE MR 750, and Philips Achieva and Ingenia models, across 21 research sites with harmonized scanning protocols to ensure data consistency. T1-weighted (T1w) images were acquired using a 3D sequence with 1mm isotropic resolution, corrected for gradient nonlinearity distortions using scanner-specific nonlinear transformations. Cortical reconstruction and volumetric segmentation were performed using FreeSurfer v7.1.1 (29). The current study considered cortical features (i.e., cortical thickness, cortical volume, cortical surface area), with cortical regions labelled according to the Destrieux atlas-based classification (mapped to 74 cortical regions per hemisphere; a total of 148 regions) (30).

### Behavioral measures

The parent-report CBCL 6-18 was used to measure the behavioral syndrome of adolescents aged 9-10 (26). The CBCL has 113 items divided into syndrome scales including Withdrawn/Depressed, Somatic Complaints, Anxious/Depressed, Rule-Breaking Behavior, Social Problems, Thought Problems, Attention Problems, and Aggressive Behavior. All syndrome scales have a T-score mean of 50 with a standard deviation of 10, with norms adjusted for gender and age groups. Scores on each syndrome scale were dichotomized as either ‘at risk’ (*T*−*score ≥* 65)or ‘low risk’ (*T*−*score* < 65)using the ‘borderline’ T-score threshold (31). Score distributions and proportions of individuals considered at risk are presented in Figure S2.

### Psychiatric diagnoses

The computerized parent version of the Kiddie Schedule for Affective Disorders and Schizophrenia (KSADS), based on DSM-5 criteria, was used to establish psychiatric diagnoses. We considered lifetime diagnoses for depression, bipolar disorder, anxiety, disruptive mood dysregulation disorder, psychosis, obsessive-compulsive disorder, post-traumatic stress disorder, and attention deficit hyperactivity disorder (ADHD). Following the protocol detailed in Duffy et al. (32), we categorized individuals into larger diagnostic classes based on specific diagnoses within each class. An Autism Spectrum Disorder (ASD) diagnosis was assigned if a parent reported a previous diagnosis.

### Latent class analysis

LCA was used to identify latent, or unobserved, subgroups within samples with homogeneities in patterns of responses to observed variables (21). We performed LCA based on the eight CBCL syndrome scales using the R package poLCA (33). We iteratively fitted LCA models from two classes to six classes and selected the final model based on the Bayesian Information Criterion (BIC) (34), which is considered a more reliable fit statistic than others (21,35). The selected LCA model was used to assign each individual to the class with the highest posterior probability. The model’s uncertainty in assigning a class membership was measured with Shannon’s entropy (*E*) (36). We set a constraint on classification uncertainty, only considering samples assigned to latent classes with relatively high certainty (*E* ≤ 0.2), to obtain psychopathological cohorts of high symptomatic homogeneity.

### Normative modelling

A conditional Variational Autoencoder (cVAE) (37) was used to develop normative models. The cVAE, a variant of the variational autoencoder, aims to learn a probabilistic mapping from the input space *X* to a latent space *Z*, from which point estimates can be decoded back to the input space *X* with minimal reconstruction error (23). The cVAE allows the encoding of the latent space and the decoding for reconstruction to be conditioned on confounding covariates, which helps avoid the need to regress out covariate effects from inputs, which is typically achieved using linear models and may remove unknown sources of variances (38). We had our models conditioned on biological sex (i.e., male/female). Technically, cVAEs are trained to maximize the following evidence lower bound (ELBO) of the marginal log-likelihood of the observed data x, conditioned on the conditional variable c:

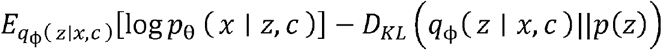

where the approximate posterior distribution q_ϕ_(*z* | *x*,*c*) and the conditional likelihood distribution p_θ_ *(z* | *z*,*c*) are modelled by the encoder and decoder with parameters ϕ and θ, respectively. The first term, therefore, forces the model to reduce reconstruction errors, while the second term forces the encoded latent distribution q (z | *x*,*c*) to be close to the prior distribution p(*z*), which is chosen as a Gaussian distribution suitable for normative modeling. A cVAE was trained for each of the cortical modalities (i.e., cortical thickness, volume, and surface area), resulting in a total of three normative models. Further technical details on variational autoencoder, model configuration and training details (including data pre-processing), are provided in the Supplement and demonstrated in Figure 1.

**Figure 1:**
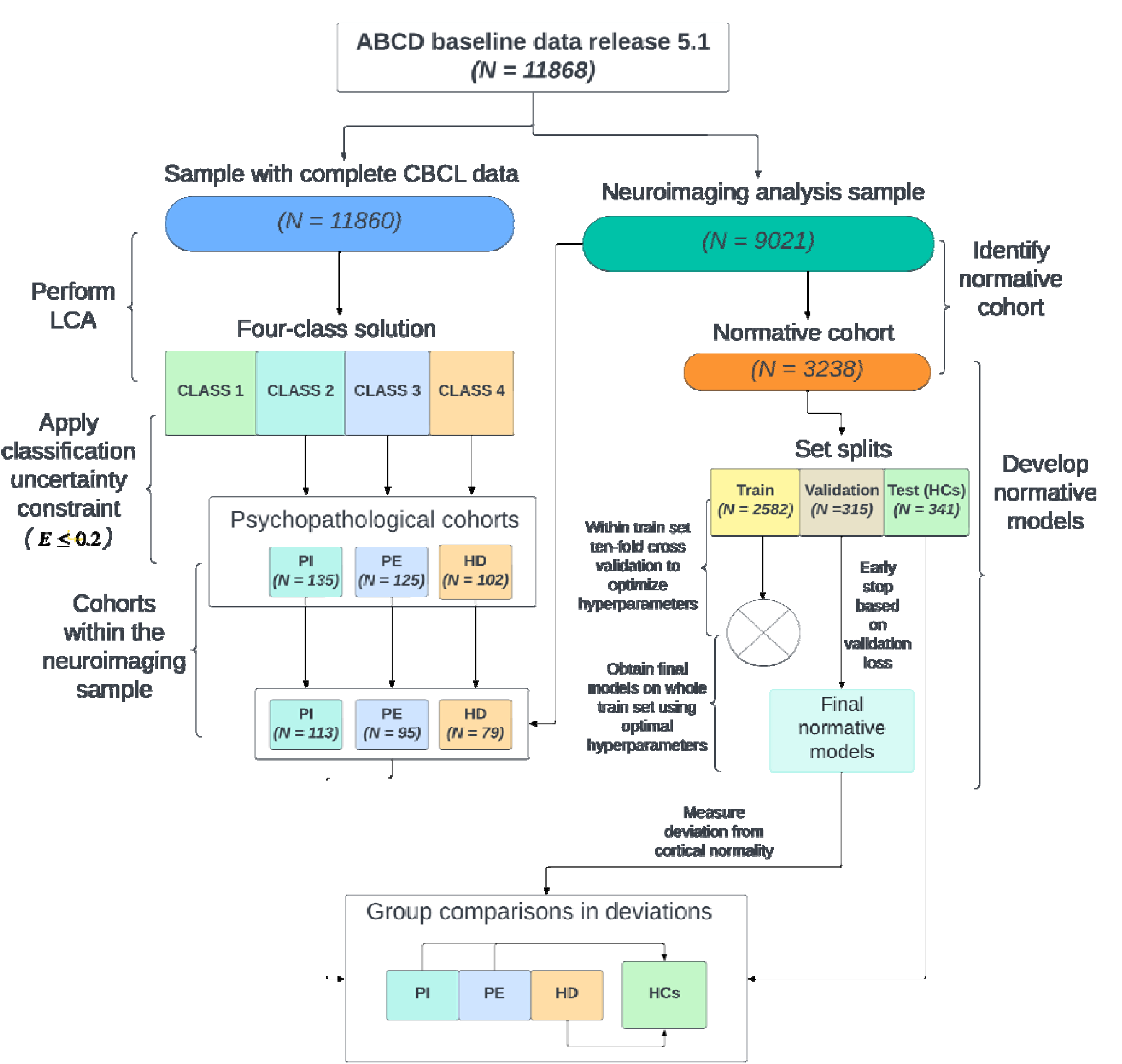
Analysis pipeline. The left side of the figure demonstrates the analysis flow of applying latent class analysis to samples with complete data for the CBCL syndrome scales from the ABCD baseline data. By applying a constraint on classification uncertainty, three psychopathological cohorts were identified. The right side of the figure demonstrates the process of developing normative models using a neuroimaging subsample drawn from the ABCD baseline data. A normative cohort was identified from the neuroimaging sample and was split into three independent sets. A 10-fold cross-validation was performed using the training set to tune a set of hyperparameters for model training. Once the optimal hyperparameters were obtained, final models were obtained using the whole training set and the training was early stopped using a separate validation set to avoid overfitting. The samples in the identified psychopathological cohorts that were present in the neuroimaging sample were compared against the testing set (i.e., HCs) to derived cortical abnormalities to specific psychopathologies.

### Normative cohort

We identified a normative cohort from our neuroimaging analysis sample as subjects without any current or past psychiatric diagnoses (including other psychiatric diagnoses not specified above) and without clinically significant behavioral syndromes (*T*−*score ≥* 65) for any of the CBCL scales, resulting in a total of *N* = 3238 subjects. The cohort was divided into Training, Validation, and Healthy Controls (HCs) (i.e., Testing set) subsamples randomly sampled from data partitions stratified by sex and household income, using an 8:1:1 split ratio. Distributions of demographic variables for each subset are shown in Figure S1.

### Deviation metrics

The metric to quantify the deviation of the whole cortex from normality was derived as the mean square error between the reconstructed feature values and the input feature values (39), as in the below equation:

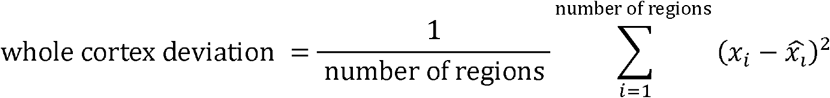

where *x*_*i*_ denotes the input value of the *i*-th cortical region and 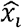 denotes the autoencoder reconstructed value of the *i*-th cortical region. The assumption behind the metric is that since the cVAE was trained to only encode healthy structures of the cortex, higher reconstruction errors (i.e., deviations) will be observed from processing cortical features of individuals with psychopathologies, whose cortical structures are hypothesized to alter pathologically (39).

The whole cortex deviation metric averages the deviations observed at individual cortical regions. A region-level deviation metric, therefore, allows an investigation of which individual regions tend to exhibit more abnormalities:

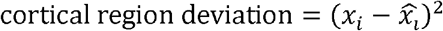

where a region level deviation is measured at the *i*-th cortical region.

To account for reconstruction errors due to an imperfect model fit, we used Cliff’s delta (δ_Cliff_(a non-parametric effect size metric (40)) to index the degree of abnormality for each region by comparing distribution differences in deviations between psychopathological cohorts and a cohort of neurotypical individuals (i.e., HCs, described above). Unlike traditional case-control studies, we note that our method measures effect size based on *deviations* quantified from normative models, instead of feature values themselves. We further assessed the robustness of effect sizes by bootstrapping the training set 1,000 times, creating simulated sets of the same size, and training multiple normative models from these sets to derive bootstrapped confidence intervals (CIs) of effect size for each cortical feature. We consider a cortical feature to exhibit statistically significant abnormality if the lower bound of the 99% CI exceeds 0.

### Sensitivity analyses

An alternative normative modeling analysis was performed comparing deviation distributions of the original latent classes defined by LCA (latent classes 2-4 without applying a constraint on classification uncertainty) against HCs. The results are reported in the Supplement.

## Results

### Latent class analysis: psychopathological cohorts

Fit statistics for all latent class solutions and a detailed descriptive analysis of the selected solution, including its individual-level classification uncertainty, comparisons between and after an exclusion of high-entropy subjects, are provided in the Supplement (Table S4-5, Figures S2-6). We selected the four-class solution according to the lowest BIC scores. By applying a constraint on classification uncertainty, we identified three psychopathological cohorts characterized by homogeneous and differential patterns of behavioral syndromes, conceptually close to broad psychopathological dimensions.

Specifically, a cohort with increased internalizing syndromes (i.e., Anxious/Depressed, Withdrawn/Depressed, and Somatic), and a relatively high prevalence of thought problems, was identified and thus labeled the *Predominantly Internalizing* cohort. A second cohort exhibited increased externalizing syndromes (i.e., Rule-Break and Aggressive) with a relatively high prevalence of attention problems and was thus labeled the *Predominantly Externalizing* cohort. Lastly, a cohort characterized by elevations of all syndromes was labeled the *Highly Dysregulated* cohort.

Since some subjects within the psychopathological cohort were not present in the neuroimaging analysis sample due to the exclusion criteria set in the *Sample* Section, they were removed from the normative modeling analysis. Table 1 summarizes the sample characteristics of the remaining sample for each psychopathological cohort against the HCs (demographic information of the original samples in each cohort is provided in Table S5). Figure 2 presents the proportions of remaining samples with clinically significant syndromes, and psychiatric diagnoses, in the cohorts.

**Table 1:** Sample characteristics of the psychopathological cohorts and the HCs.

|  | Healthy controls | Predominantly internalizing | Predominantly externalizing | Highly dysregulated |
| --- | --- | --- | --- | --- |
| N | 341 | 113 | 95 | 79 |
| **Biological sex, n (%)** |  |  |  |  |
| Male | 162 (47.5%) | 53 (46.9%) | 57 (60.0%) | 53 (67.1%) |
| Female | 179 (52.5%) | 60 (53.1%) | 38 (40.0%) | 26 (32.9%) |
| **Age in months** |  |  |  |  |
| mean (SD) | 118.83 (7.22) | 119.44 (7.11) | 119.16 (7.41) | 118.94 (7.17) |
| **Race/ethnicity, n (%)** |  |  |  |  |
| White | 177 (51.9%) | 55 (48.7%) | 36 (37.9%) | 32 (40.5%) |
| Black | 52 (15.2%) | 7 (6.2%) | 31 (32.6%) | 16 (20.3%) |
| Hispanic | 70 (20.5%) | 35 (31.0%) | 9 (9.5%) | 20 (25.3%) |
| Asian | 12 (3.5%) | 1 (0.9%) | 0 (0.0%) | 0 (0.0%) |
| Other | 30 (8.8%) | 15 (13.3%) | 19 (20.0%) | 11 (13.9%) |
| **Annual income in \$, n (%)** | | | | |
| < 5,000 | 12 (3.5%) | 1 (0.9%) | 10 (10.5%) | 12 (15.2%) |
| 5,000 - 11,999 | 12 (3.5%) | 3 (2.7%) | 11 (11.6%) | 9 (11.4%) |
| 12,000 - 15,999 | 9 (2.6%) | 7 (6.2%) | 4 (4.2%) | 3 (3.8%) |
| 16,000 - 24,999 | 15 (4.4%) | 4 (3.5%) | 5 (5.3%) | 7 (8.9%) |
| 25,000 - 34,999 | 18 (5.3%) | 14 (12.4%) | 12 (12.6%) | 7 (8.9%) |
| 35,000 - 49,999 | 26 (7.6%) | 18 (15.9%) | 6 (6.3%) | 7 (8.9%) |
| 50,000 - 74,999 | 39 (11.4%) | 16 (14.2%) | 12 (12.6%) | 10 (12.7%) |
| 75,000 - 99,999 | 45 (13.2%) | 10 (8.8%) | 4 (4.2%) | 4 (5.1%) |
| 100,000 - 199,999 | 94 (27.6%) | 26 (23.0%) | 19 (20.0%) | 7 (8.9%) |
| > 200,000 | 40 (11.7%) | 7 (6.2%) | 3 (3.2%) | 3 (3.8%) |
| Not Provided | 31 (9.1%) | 7 (6.2%) | 9 (9.5%) | 10 (12.7%) |

**Figure 2:**
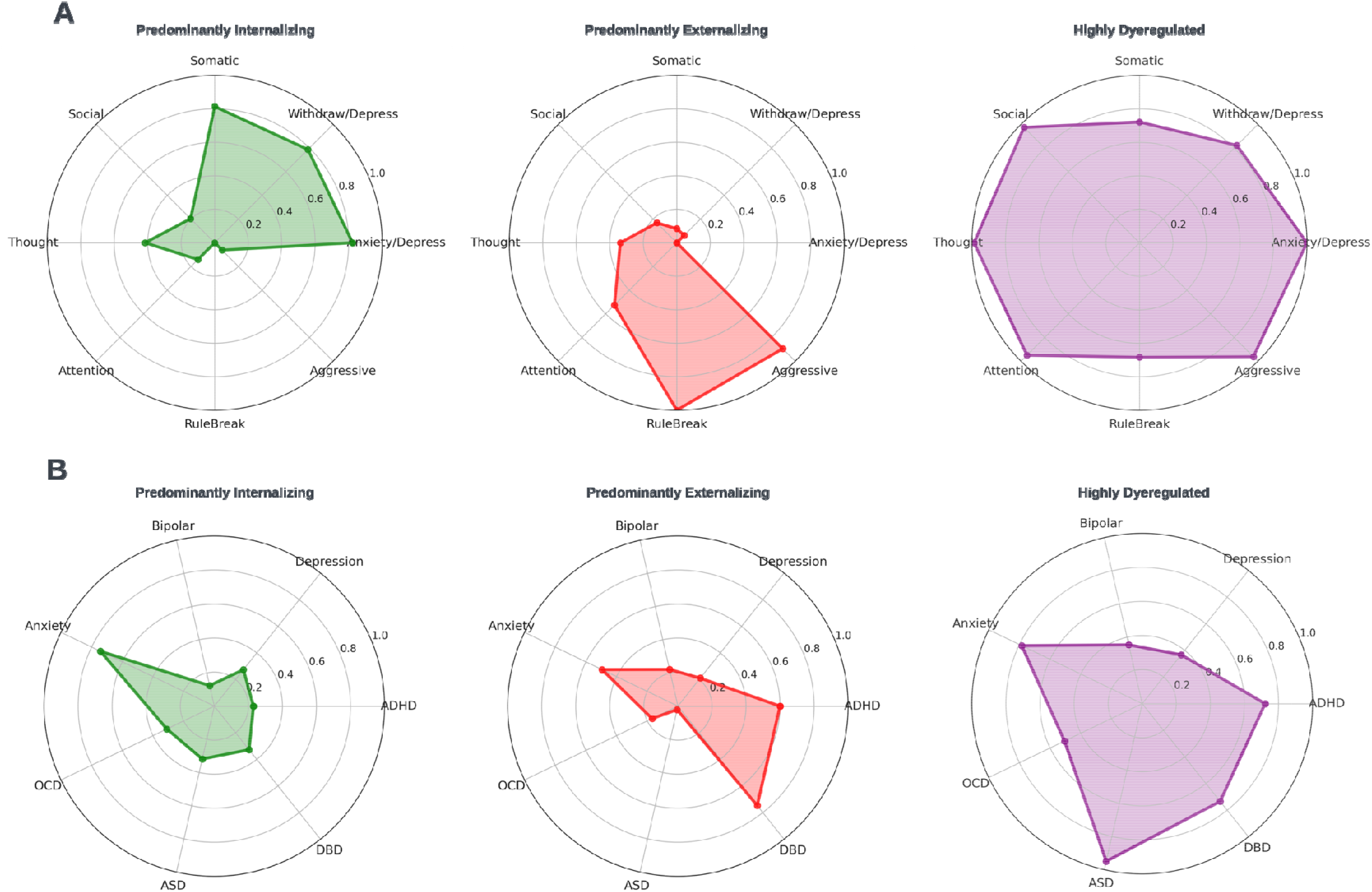
Radar charts. A – A set of three radar charts demonstrating the proportions of subjects within each of the psychopathological cohorts having clinically significant syndromes across the CBCL syndrome scales. B – A set of three radar charts demonstrating the proportions of subjects within each of the psychopathological cohorts having psychiatric diagnoses.

### Normative modelling: whole cortex deviation

Figure 3A presents the distributions of the whole cortex deviation for each psychopathological cohort compared against the HCs. Given the Shapiro-Wilk tests indicated that none of these deviation scores were normally distributed (p < .05; see test results in Table S6), Mann-Whitney U tests (two-sided) were performed to determine statistically significant differences between groups. A false discovery rate (FDR) correction was applied for multiple tests within each modality. We observed a significant difference in cortical thickness for the *Predominantly externalizing* cohort (*FDR-corrected, p* < . 05, δ_Cliff_ = 0.256) and the *Highly Dysregulated* cohort (*FDR-corrected, p* < .05, δ_Cliff_ = 0.222). All other comparisons did not reach statistical significance, showing no significant difference in deviations at a whole cortex level.

**Figure 3:**
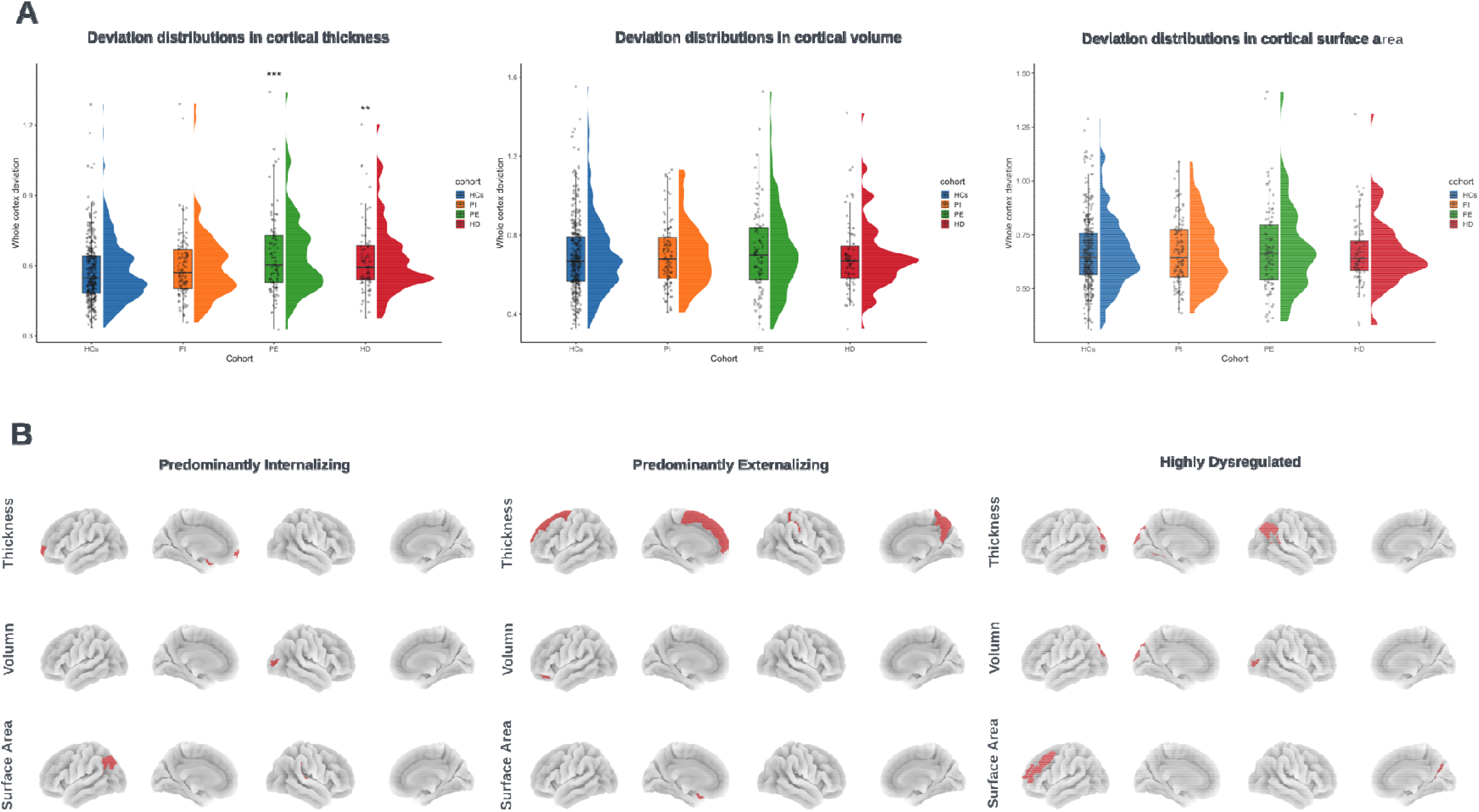
Cortical abnormalities. A – Comparisons in whole cortex deviation distributions of the psychopathological cohorts against the HCs in terms of cortical thickness, volume, surface area. Within each modality, Mann-Whitney U tests with Benjamini–Hochberg FDR correction (correcting for three comparisons tests within each modality) assessed statistical significance: * for FDR corrected at p < 0.05, ** for *FDR corrected at p* < 0.01, *** for *FDR corrected at p* < 0.001. B – Region specific cortical abnormalities in terms of cortical thickness, volume, surface area for each of the psychopathological cohort. Only regions exhibiting significantly, and meaningfully, higher deviations (mean δ_Cliff_ ≥ 0.15) are plotted in red.

### Normative modeling: individual region deviation

CIs of effect sizes (δ_Cliff_) for each significant cortical feature are presented in Figure S10-12. Here, we report the significant features with at least a small to medium effect size, indicated by a mean bootstrapped effect size larger than or equal to 0.15 (mean δ_Cliff_ ≥ 0.15) (41). Figure 3B shows the spatial representation of these features for each cohort.

Although no significant difference in deviations was observed at a whole cortex level, an analysis of deviations at individual cortical regions revealed regions at which the *Predominantly internalizing* cohort tends to exhibit significantly more deviations than HC, including the left transverse frontopolar gyri/sulci and left planum polare of the superior temporal gyrus in cortical thickness, right middle occipital sulcus and lunatus sulcus in cortical volume, and the left angular gyrus and the right planum temporale in cortical surface area.

In alignment with the significant deviation observed at a whole cortex level, the *Predominantly externalizing* cohort exhibited significantly more deviations at multiple regions in cortical thickness, including the left superior frontal gyrus, left superior frontal sulcus, left vertical ramus of the anterior segment of the lateral sulcus, right precuneus, and right postcentral sulcus. A few other regions also show significantly more deviations in other modalities, including the left orbital sulci in cortical volume, and the left planum polare of the superior temporal gyrus in cortical surface area.

Similarly, multiple regions exhibited significantly more deviations in cortical thickness for the *Highly Dysregulated* cohort including the left superior occipital gyrus, left middle occipital sulcus and lunatus sulcus, left medial occipito-temporal sulcus and lingual sulcus, and the right angular gyrus. A few regions showed significantly more deviations including the left superior occipital gyrus, and the right middle occipital sulcus and lunatus sulcus in cortical volume, the left middle frontal gyrus and the right transverse temporal sulcus in cortical surface area.

The sensitivity analysis compared the original latent classes of high probabilities of at-risk behavioral syndromes defined by the four-class LCA solution (latent class 2-4, see Figure S3) against HCs. One cortical region (left orbital sulci) showed significantly more deviations in cortical volume for the latent class 3 with a small mean effect size, but all other cortical features for each latent class had small, negligible mean effect sizes (mean δ_Cliff_ < 0.15). Bootstrapped CIs of these effect sizes are shown in Figure S14-16.

## Discussion

The current study applied LCA to empirically identify psychopathological cohorts of homogeneous syndrome patterns in a large cohort of preadolescents in the ABCD study. By applying an additional constraint on classification uncertainty, we obtained three psychopathological cohorts, labelled *Predominantly internalizing, Predominantly externalizing*, and *Highly Dysregulated*, each characterized by differential elevations of behavioral syndromes clustered clearly into the CBCL internalizing syndrome scales, the externalizing syndrome scales, and all syndrome scales, respectively, thereby representing the clinical end of broad psychopathological dimensions (i.e., internalizing/externalizing, *p-factor*). Due to their high within-cohort symptomatic homogeneities, they served relatively precise symptomatic targets to allow higher sensitivity in identifying syndrome-specific neurobiological abnormalities.

By comparing the psychopathological cohorts against a group of neurotypical individuals in deviations from the normative patterns of cortical structures, our study identified cortical thickness anomalies as relevant to psychopathologies in preadolescents in the ABCD data, particularly regarding the externalizing syndromes. Notably, this finding contradicts previous association studies using the same dataset (i.e., ABCD baseline data) (17–19), which reported robust associations between psychopathologies and cortical volume and surface area, but not cortical thickness.

A potential explanation for this contradiction in findings lies in methodological differences. Our normative modeling approach identifies deviations from the normality of cortical thickness, disregarding their directions. In other words, our method captures abnormalities indicative of psychopathologies whether they manifest as abnormal cortical thinning or thickening relative to normal developments seen in neurotypical individuals, thereby providing a more flexible, hypothesis-free, view of structural abnormalities associated with psychopathologies. In contrast, previous studies using a dimensional approach to psychopathologies focus on directional associations between neurological correlates and psychopathologies, operationalized as continuous variables (e.g., factor scores from factor analysis) (17,18,19,41,42). These association studies impose a directional assumption that precludes the possibility of abnormalities in cortical thickness manifested in different directions across individuals with phenotypically similar psychopathologies. Previous association studies, therefore, may suggest a lack of linear, or monotonic, association between cortical thickness and psychopathologies, as opposed to *any* relationships.

Furthermore, our analysis revealed relatively distinct patterns of cortical abnormalities to divergent dimensional psychopathologies at a group level. Specifically, we found localized abnormalities regarding cortical thickness in the left frontopolar cortex (FPC) associated with the *Predominantly internalizing* cohort. Abnormality in this region may complement a structural understanding of previous functional observations on internalizing disorders in adult samples, which report, for instance, reduced functional connectivity of the left FPC to the right lateral orbitofrontal cortex, associated with increased depressive symptoms (44). Furthermore, we found the cohort to exhibit asymmetric abnormalities in the superior temporal gyrus (i.e., left planum polare and right planum temporale), a region implicated in positive symptoms (e.g., hallucinations, thought disturbances) seen in schizophrenia, potentially reflecting the thought problems in the cohort.

The *Predominantly externalizing* cohort, in contrast, showed more frequent abnormalities in the left superior frontal cortex (SFC), left Orbitofrontal Cortex (OFC), and right Parietal Cortex (PC) (i.e., postcentral sulcus and Precuneus), primarily in terms of cortical thickness. Abnormalities in these regions are highly consistent with previous findings on externalizing behavior/disorders. In particular, a reduction in the left SFC thickness has been implicated in increased impulsivity, which is associated with inattention, commonly observed in ADHD (45), potentially via its functional role in response inhibition and executive attention (46,47). Similarly, OFC abnormalities, via its altered structural white matter connections with the amygdala (48), have been observed in externalizing behavior/disorders (e.g., conduct disorder) (14,49). Notably, we also identified a structural abnormality in the right PC, a region engaged in goal-directed attentional processing (50), which, together with abnormalities in the SFC and OFC, aligns with attentional dysfunction as a hallmark of externalizing syndromes.

The *Highly Dysregulated* cohort may conceptually represent the clinical end of the *p-factor* continuum (51), implicating the development of a wide range of syndromes/disorders. For this cohort, we found a concentration of structural abnormalities in the occipital cortex. This finding aligns with Romer and colleagues’ works on the *p-factor* (42,43,52), which have associated structural and functional alterations in the occipital cortex with higher *p-factor* scores. Our findings may, therefore, complement this association. Moreover, it is notable that the *Highly Dysregulated* cohort has a very high prevalence of ASD (95.05%, see Figure 2B). This may suggest the observed <u>occipital cortex abnormalities as a neurodevelopment basis of autistic traits</u> (53–55), which may mediate the development of psychopathologies in other domains (56).

Our findings should be interpreted in the context of a few methodological limitations. First, we used cohorts of preadolescents with clinically significant psychopathologies to conceptually represent the clinical end of broad psychopathological dimensions. Although such an approach may result in neurobiological correlates of clinical relevance, the subclinical range of variations in these dimensional phenotypes are ignored. Second, our identification of clinical cohorts with high symptomatic homogeneities necessitated the exclusion of large proportions of individuals (see Table S3 and Supplemental information) with clinically significant syndromes that deviate from the patterns our analysis revealed. This exclusion, therefore, limits the generalizability of our findings to other clinical cohorts (e.g., individuals with only attention problems). Third, although our squared, non-directional, measurement of deviation from normality may increase sensitivity for identifying abnormalities, it simultaneously precludes a directional interpretation both at a group and individual level (e.g., abnormal cortical thickening or thinning). Finally, it is important to note that our cross-sectional analysis can only reveal associative brain-behavioral relations, which precludes any causal interpretations.

In summary, the current study uses psychopathological cohorts of high symptomatic homogeneity to represent the high-risk end of broad psychopathological dimensions, and revealed distinct patterns of cortical abnormalities between them. We also found that using a normative approach identifies cortical thickness abnormalities as related to psychopathologies in the ABCD data, especially those featuring externalizing. Our study underscores the value of a person-centred analytic approach (e.g., LCA) to complement variable-centred analysis (e.g., factor analysis) in the study of dimensional psychopathologies. When combined with normative modeling (e.g., autoencoder-based normative models), this approach allows group-level comparisons in deviations from normality, quantified from large non-clinical populations, as an alternative to an associational methodology to offer new insights into the neurobiological correlates of psychopathologies.

