## Supplementary material for "Normative models reveal distinct cortical abnormalities to dimensions of psychopathologies in preadolescents"

Qingkun Deng et al.

**Supplementary materials and methods**

**Variational autoencoder**

**Technical details.** The Variational autoencoder (VAE) aims to learn a probabilistic mapping from the input space X to a latent space Z, from which point estimates can be decoded back to the input space X with minimal reconstruction error (1). Technically, VAEs are trained to maximize the following evidence lower bound (ELBO) of the marginal log-likelihood of the observed data $x$:

$$E_{q_{\phi}\left( z \mid x \right)}\left[ \log p_{\theta}\left( x \mid z \right) \right]-D_{KL}\left( q_{\phi}\left( z \mid x \right)||p\left( z \right) \right)$$

The ELBO consists of two terms: the expected log-likelihood of the data given the latent variables $z$, and the KL divergence between the approximate posterior $q\left( z \mid x,c \right)$ and the prior distribution $p\left( z \right)$. The first term forces the reconstruction error between the input values and the reconstructed values to be reduced and the second forces the approximate posterior to be close to the prior distribution, set to a Gaussian.

However, computing the gradient of the ELBO with respect to the parameters $\phi$ of the encoder is intractable. For a Gaussian encoder $q_{\phi}\left( z \mid x \right)\mathcal{=N}\left( z \mid f_{e}^{\mu}\left( x \right),f_{e}^{\sigma}\left( x \right) \right)$, where the encoder $f_{e}$ outputs both the mean $\mu$ and variance $\sigma$ of the latent variable $z$, the reparameterization trick can be used to sample from $z$ and thereby enable the calculation of gradients with respect to $\phi$ (2).

In our study, we applied conditional VAE (cVAE), which is a variant of VAEs that allows the encoding of the latent space and the decoding for reconstruction to be conditioned on confounding covariates. cVAEs are trained to maximize the following updated evidence lower bound (ELBO) of the marginal log-likelihood of the observed data $x$ conditioned on the conditional variable $c$:

$$E_{q_{\phi}\left( z \mid x,c \right)}\left[ \log p_{\theta}\left( x \mid z,c \right) \right]-D_{KL}\left( q_{\phi}\left( z \mid x,c \right)||p\left( z \right) \right)$$

The confounding effects of the covariates can therefore by removed by conditioning the encoding and decoding distributions on the levels of the covariates. We consider biological sex (i.e., male/female) as the conditional variable, to avoid linearly regressing its effects out from the cortical features beforehand.

**Model training.** The normative cohort was divided into three sets (i.e., train, validation, and Healthy Controls (HCs), as detailed in the main paper). The distributions of demographic variables of each subset are shown in Figure S1. Before model training, the raw cortical features were pre-processed. To avoid deviation metrics being biased by confounders, we first applied the NeuroCombat algorithm (3) to remove the site effect from the cortical features, while including biological sex and interview age as covariates. Although the site effects were estimated using all sets together (i.e., Train/Validation/HCs), thus a `leaky site correction' method (4), it has been observed to have little or no effect on prediction performance on out-of-sample data (4). Subsequently, we removed the effects of age (i.e., interview age in months) and intracranial volume from the training sets by fitting a multivariate linear regression and extracting the residuals. Then, we applied the regression models fitted to the training data to the other sets and extracted the residuals. To account for the effect of biological sex, we created one-hot encoded vectors with two positions to represent the sample's biological sex, which were fed with the cortical features to the cVAEs such that the model distributions were conditioned on biological sex. Data standardization was performed independently for each cortical feature of the training set to have a mean of 0 and a Standard Deviation (SD) of 1. The relevant statistics (mean and SD) from the training set were used to scale the corresponding cortical features for the other sets. We retained the final model parameters at the epoch where the model loss was the lowest on the validation set after the validation loss stopped improving for 50 epochs.

**Model configurations.** We performed a grid search via a ten-fold cross-validation of the training set to obtain an optimal set of hyperparameters, which resulted in the lowest average cross-validation loss, for each of the cortical modalities (i.e., cortical thickness, volume, and surface area). The optimal hyperparameter sets for each modality are shown in Table S1. We opted not to tune dimensionality size, as increasing latent dimensions typically reduces validation loss but at the expense of model complexity. Also, the impact of dimensionality size on deviation measures in normative models is not well understood, and optimizing for validation loss may not enhance the relevance of deviation metrics for psychopathologies. We, therefore, fixed the dimensionality size at 10 for all models. The RELU non-linear activation function was applied to each linear layer to allow the model to learn non-linear relationships.

Table S1. Hyperparameter search space. For hidden dimension, two values within a list refer to two linear layers of multiple nodes (e.g., [50, 50] refers to two linear layers of 50 nodes).

| **Hyperparameter** | **Search space** |
| --- | --- |
| Batch Size | [64, 128, 256] |
| Learning Rate | [0.01, 0.005, 0.001, 0.0005] |
| Latent Dimension | [10] |
| Hidden Dimension | [30], [30, 30], [40], [40, 40], [50], [50, 50] |

Table S2. Optimal hyperparameter set for each cortical modality.

| **Feature Type** | **Batch Size** | **Learning Rate** | **Latent Dimension** | **Hidden Dimension** |
| --- | --- | --- | --- | --- |
| Cortical Thickness | 256 | 0.0005 | 10 | 40 |
| Cortical Volume | 256 | 0.005 | 10 | 30 |
| Cortical Surface Area | 256 | 0.0005 | 10 | [30, 30] |

Figure S1. Distributions of demographic variables for the train/validation/HCs subsets of the normative cohort.

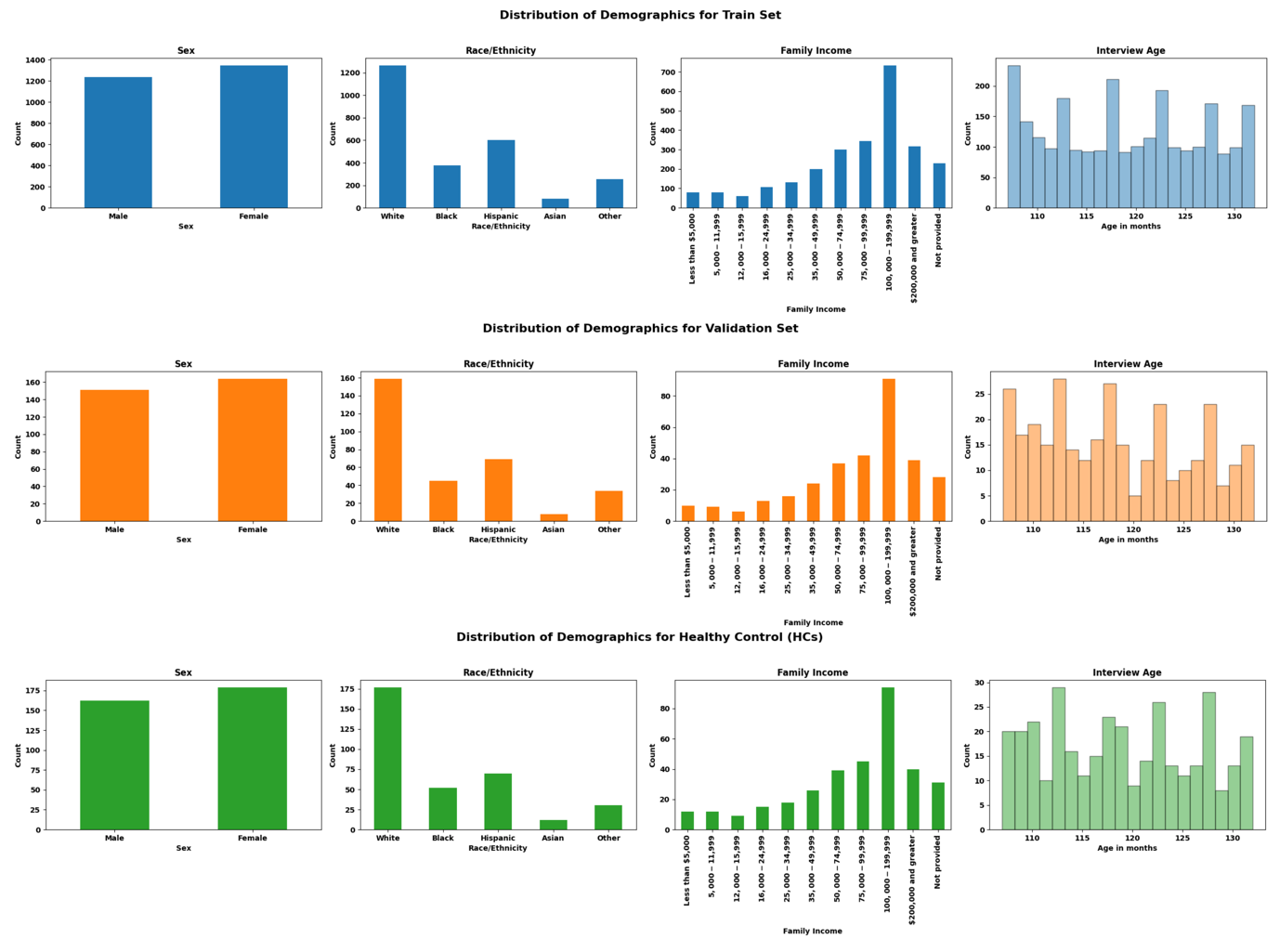

**Supplementary results**

**Latent class analysis**

**Sample characteristics.** Sample characteristics of the sample with complete Child Behavior Checklist (CBCL) data for performing latent class analysis (LCA) are summarized in Table S3. Distributions of each CBCL syndrome scale score are shown in Figure 2.

**Fit statistics.** Table S4 summarizes the fit statistics for class solutions from 2 classes to 6 classes. The BIC, which is generally considered to be the most reliable fit statistic for LCA (5,6), suggested a four-class solution. The adjusted BIC indicated a five-class solution, whereas the consistent AIC indicated a four-class solution. Both the LMR test and AIC support add up to six classes. Despite the disagreement among the fit statistics, we selected the four-class solution based on the BIC, which had an acceptable entropy of 0.90, suggesting a relatively accurate definition of classes.

**Latent class sample characteristics.** Figure S3 presents the probabilities of having at-risk behavioral syndromes ($T-score \geq65$) across the CBCL syndrome scales for each of the latent classes defined in the four-class solution. Except for the latent class 1, which had low probabilities of being above the risk threshold, the other three latent classes (2-4) had relatively elevated probabilities of at-risk behavioral syndromes. The sample characteristics for the four latent classes are summarized in Table S5.

**Individual classification uncertainty.** The four-class solution assigns each subject to the four latent classes based on the class with the highest posterior probability. Therefore, some subjects were classified with high certainty while others were not. High levels of classification uncertainty within the latent classes may result in high levels of symptomatic variability or heterogeneity within classes. Even though the relative entropy for the four-class solution was high (0.9), suggesting an overall acceptable precision in LCA classification, we further quantified the uncertainties of the model in classifying each subject using Shannon’s entropy ($E$).

Figure S4 demonstrates the distributions of Shannon's entropy for every subject within each latent class. Relative to the other latent classes, class 1 had low levels of classification uncertainties, indicated by a mean $E$ of 0.2 ($Standard Deviation, SD = 0.15$). The other three classes had relatively high uncertainties ($mean E = 0.43, SD = 0.23$ for class 2, $mean E = 0.35, SD = 0.27$ for class 3 and $mean E = 0.35, SD = 0.27$ for class 4). This analysis of individual-level Shannon’s entropy, therefore, revealed classification uncertainties within the latent classes of relatively high probabilities of at-risk behavioral syndromes (class 2-4).

These classification uncertainties are reflected in the relatively undefined syndrome patterns of classes 2-4, shown in Figure S5. Figure 6 shows the variation ($\text{Mean}\pm1 \text{Standard Deviation}$) of each syndrome scale for these latent classes. Of note, latent class 2 exhibits the most varied symptomatic patterns, as reflected by mean scores for its characteristic internalizing syndrome scales (i.e., anxiety/depress, withdraw/depress, somatic) falling below the risk threshold of 65, together with relatively large variation, suggesting within-class symptomatic heterogeneities.

**Comparisons between and after subject exclusion.** In our study, we applied an additional constraint on classification uncertainty ($E\leq0.2$) in order to obtain psychopathological cohorts with relatively high symptomatic homogeneity within each class. Figure S7-8 demonstrates the change in syndrome patterns before and after an exclusion of subjects with relatively high Shannon’s entropy ($E>0.2$). A total of, $N = 711$, $N = 167$, $N = 140$, samples were excluded from classes 2, 3, and 4, respectively. After the exclusion, relatively characteristic and homogeneous syndrome patterns emerged for classes 2-4, shown in Figure S9, which are therefore labeled the *Predominantly internalizing, Predominantly externalizing,* and *Highly dysregulated* cohort. The demographics of these cohorts are shown in Table S6.

**Normative modelling**

**Shapiro-Wilk tests.** Shapiro-Wilk tests indicated that none of the whole cortex deviation scores were normally distributed ($p < .05$). We, therefore, performed Mann-Whitney U tests. Shapiro-Wilk test results are summarized in Table S6.

**Cortical region deviation.** Figure S10-12 presents the bootstrapped confidence intervals of effect size of group differences in deviation scores for each cortical feature for each psychopathological cohort.

**Sensitivity analysis**

A comparative analysis was performed to compare the original latent classes of high probabilities of developing clinically significant symptoms (i.e., class 2-4) (before applying constraint on classification uncertainty) against HCs in deviation distributions. A total of, $N = 663$, $N = 224$, $N = 187$, subjects within the latent class 2, 3, and 4, were available in the neuroimaging sample and were thus included in the analysis. Overall, the results showed a notable decrease in effect sizes for all individual cortical features and less interpretable patterns of cortical abnormalities.

**Whole cortex deviation.** Figure S13 shows the distributions of the whole cortex deviation for each latent class compared against the deviation distribution of the HCs. The Mann-Whitney U tests indicated significant group differences for all latent classes in terms of cortical thickness. All other comparisons did not reach statistical significance.­

**Cortical region deviation.** As shown in Figure S14-16, except one cortical region showed significantly more deviations with a small mean effect size (left orbital sulci, $mean Cliff^{'}s delta\geq0.15$) in cortical volume for latent class 3, all other cortical features had negligible mean effect sizes. These generally smaller effect sizes are likely attributable to more varied within-class syndrome patterns prior to the exclusion of high-entropy subjects. The sensitivity analysis results, therefore, support the value of isolating cohorts of higher symptomatic homogeneity, at the expense of excluding samples, for enhanced sensitivity in identifying syndrome-specific neurological abnormalities.

Table S3. Demographics of the sample used for latent class analysis. Two subjects had missing values for Race/Ethnicity and one for age.

| **Category** | **Count** |
| --- | --- |
| **N** | 11860 |
| **Age in Months** |  |
| Mean (SD) | 118.98 (7.50) |
| **Biological Sex, n (%)** |  |
| Male | 6185 (52.15%) |
| Female | 5672 (47.82%) |
| Intersex-Male | 3 (0.03%) |
| **Race/Ethnicity, n (%)** |  |
| White | 6169 (52.02%) |
| Black | 1782 (15.03%) |
| Hispanic | 2408 (20.31%) |
| Asian | 252 (2.13%) |
| Other | 1247 (10.52%) |
| **Annual Income in Dollars, n (%)** |  |
| Less than 5,000 | 417 (3.52%) |
| 5,000 - 11,999 | 421 (3.55%) |
| 12,000 - 15,999 | 273 (2.30%) |
| 16,000 - 24,999 | 523 (4.41%) |
| 25,000 - 34,999 | 654 (5.51%) |
| 35,000 - 49,999 | 933 (7.87%) |
| 50,000 - 74,999 | 1498 (12.63%) |
| 75,000 - 99,999 | 1569 (13.23%) |
| 100,000 - 199,999 | 3309 (27.90%) |
| 200,000 and greater | 1250 (10.54%) |
| Not provided | 1013 (8.54%) |

Table S4. Fit statistics for LCA models with 2 to 6 classes. LL = log-likelihood; AIC = Akaike information criterion; BIC = Bayesian information criterion; aBIC = sample-size adjusted BIC; CAIC = consistent Akaike information criterion; LMR ($p$) = ad-hoc adjusted likelihood ratio test (LRT) test p-values.

| **Number of classes** | **AIC** | **BIC** | **aBIC** | **CAIC** | **LMR** | **Entropy** |
| --- | --- | --- | --- | --- | --- | --- |
| Class-2 | 36785.25 | 36910.73 | 36856.7 | 36927.73 | / | 0.92 |
| Class-3 | 36377.37 | 36569.27 | 36486.65 | 36595.27 | 0 | 0.86 |
| Class-4 | 36091.6 | 36349.93 | 36238.7 | 36384.93 | 0 | 0.9 |
| Class-5 | 36028.83 | 36353.59 | 36213.77 | 36397.59 | 0 | 0.92 |
| Class-6 | 36017.7 | 36408.89 | 36240.46 | 36461.89 | 0 | 0.83 |

|  | **Class 1** | **Class 2** | **Class 3** | **Class 4** |
| --- | --- | --- | --- | --- |
| **N** | 10480 | 846 | 292 | 242 |
| **Biological sex, n (%)** |  |  |  |  |
| Male | 5374 (51.3%) | 476 (56.3%) | 174 (59.6%) | 161 (66.5%) |
| Female | 5103 (48.7%) | 370 (43.7%) | 118 (40.4%) | 81 (33.5%) |
| **Age in months** |  |  |  |  |
| mean (SD) | 118.79 (7.49) | 118.83 (7.57) | 119.82 (7.69) | 118.91 (7.38) |
| **Race/ethnicity, n (%)** |  |  |  |  |
| White | 5505 (52.5%) | 443 (52.4%) | 116 (39.7%) | 105 (43.4%) |
| Black | 1552 (14.8%) | 95 (11.2%) | 86 (29.5%) | 49 (20.2%) |
| Hispanic | 2126 (20.3%) | 190 (22.5%) | 41 (14.0%) | 51 (21.1%) |
| Asian | 243 (2.3%) | 8 (0.9%) | 1 (0.3%) | 0 (0.0%) |
| Other | 1053 (10.0%) | 110 (13.0%) | 48 (16.4%) | 36 (14.9%) |
| **Annual income in $, n (%)** |  |  |  |  |
| < 5,000 | 344 (3.3%) | 27 (3.2%) | 24 (8.2%) | 22 (9.1%) |
| 5,000 - 11,999 | 333 (3.2%) | 36 (4.3%) | 29 (9.9%) | 23 (9.5%) |
| 12,000 - 15,999 | 222 (2.1%) | 27 (3.2%) | 15 (5.1%) | 9 (3.7%) |
| 16,000 - 24,999 | 435 (4.2%) | 52 (6.1%) | 19 (6.5%) | 17 (7.0%) |
| 25,000 - 34,999 | 546 (5.2%) | 59 (7.0%) | 26 (8.9%) | 23 (9.5%) |
| 35,000 - 49,999 | 787 (7.5%) | 95 (11.2%) | 24 (8.2%) | 27 (11.2%) |
| 50,000 - 74,999 | 1317 (12.6%) | 113 (13.4%) | 36 (12.3%) | 32 (13.2%) |
| 75,000 - 99,999 | 1417 (13.5%) | 112 (13.2%) | 24 (8.2%) | 16 (6.6%) |
| 100,000 - 199,999 | 3019 (28.8%) | 204 (24.1%) | 50 (17.1%) | 36 (14.9%) |
| > 200,000 | 1175 (11.2%) | 52 (6.1%) | 11 (3.8%) | 12 (5.0%) |
| Income Not Provided | 885 (8.4%) | 69 (8.2%) | 34 (11.6%) | 25 (10.3%) |

Table S5. Sample characteristics of the four latent classes in the four-class solution.

Figure S2. The average probabilities of class memberships for each of the CBCL syndrome scales.

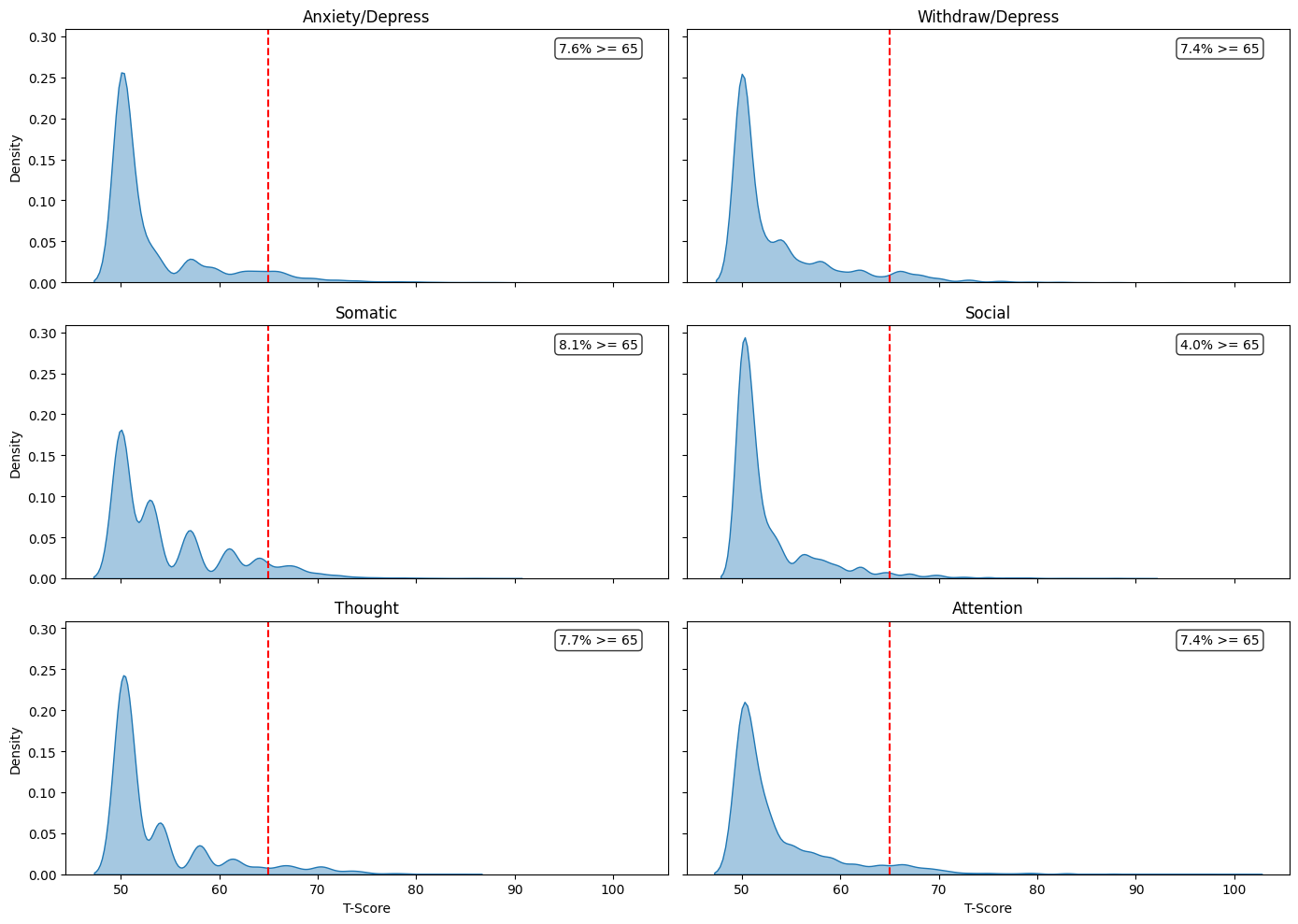

Figure S3. The average probabilities of class memberships for each of the CBCL syndrome scales. Of N = 11,860 participants, 10480 (88.36%) participants were assigned to class 1, 846 (7.13%) participants were assigned to class 2, 292 (2.46%) participants were assigned to the class 3, and 242 (2.04%) participants were assigned to class 4.

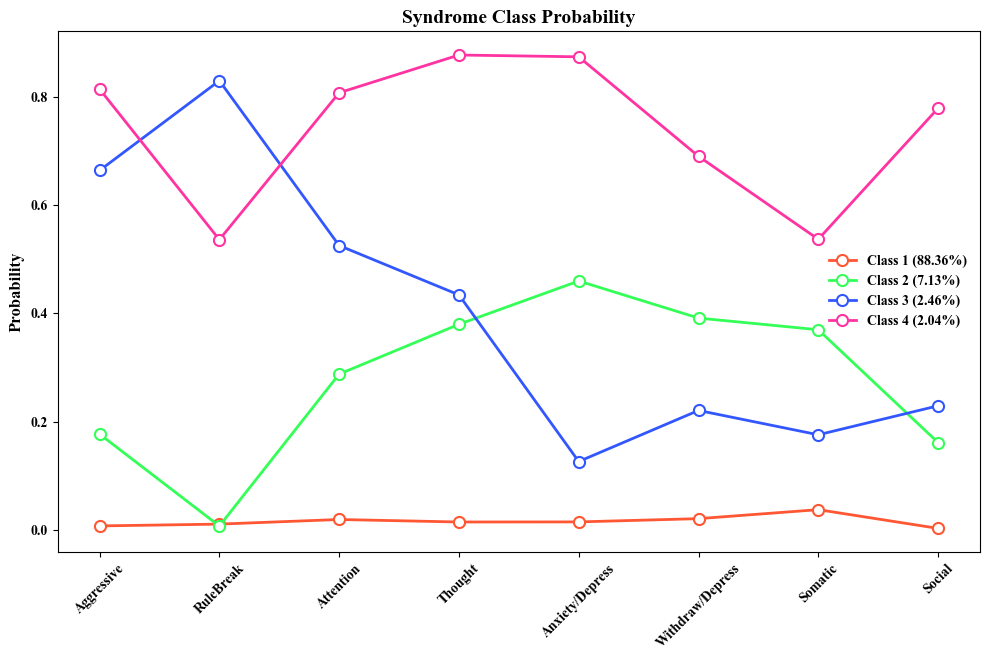

Figure S4. Distribution of Individual entropy ($E$) scores for each of the latent classes in the four-class solution.

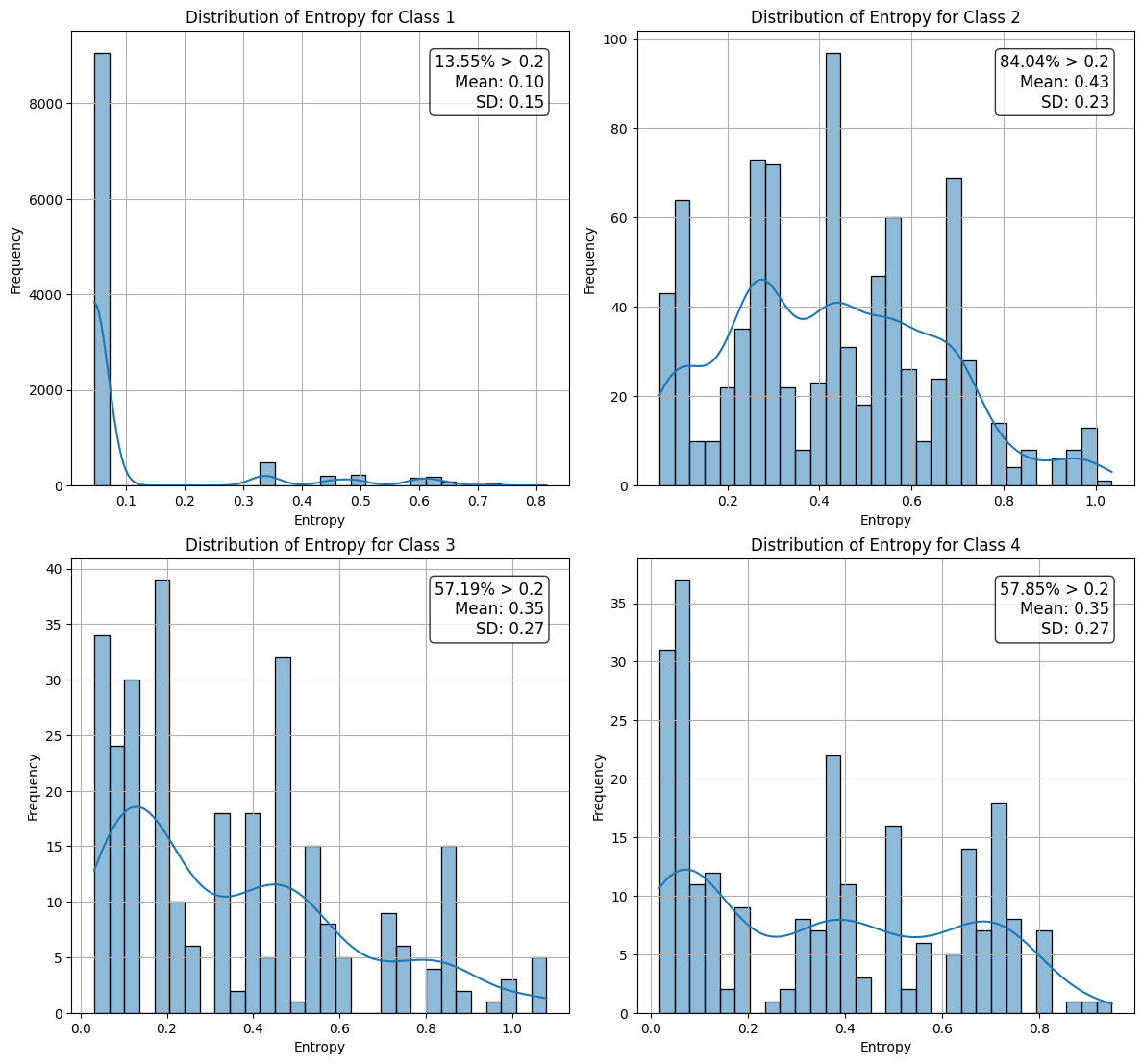

Figure S5. Subject-level syndrome patterns across the CBCL syndrome scales of class 2-4.

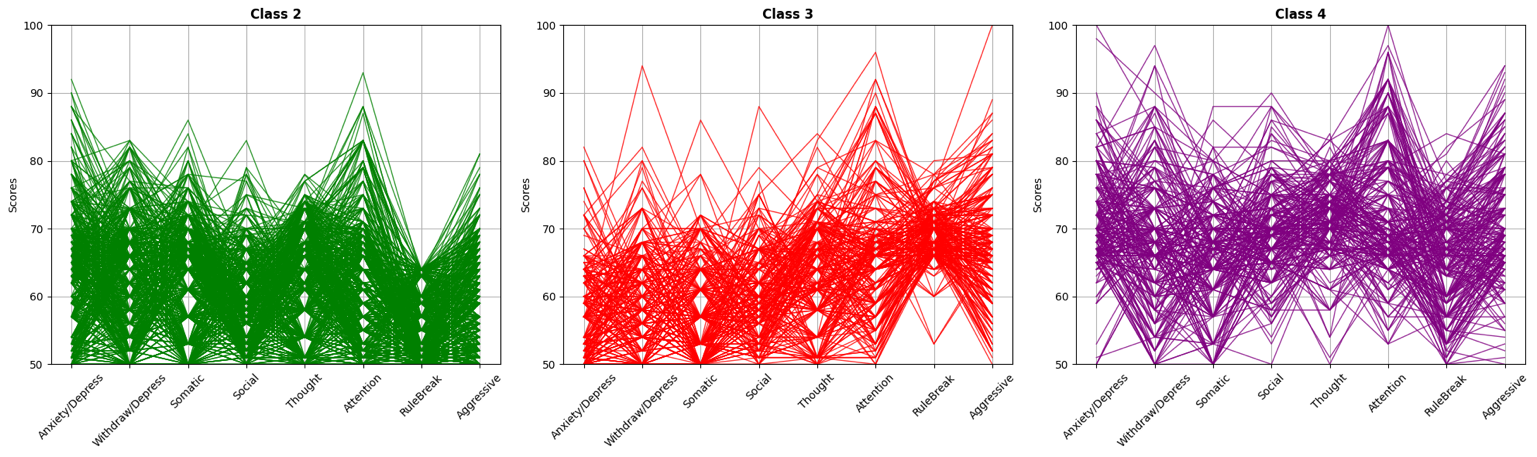

Figure S6. Mean and variation of syndrome scores across the CBCL syndrome scales of class 2-4.

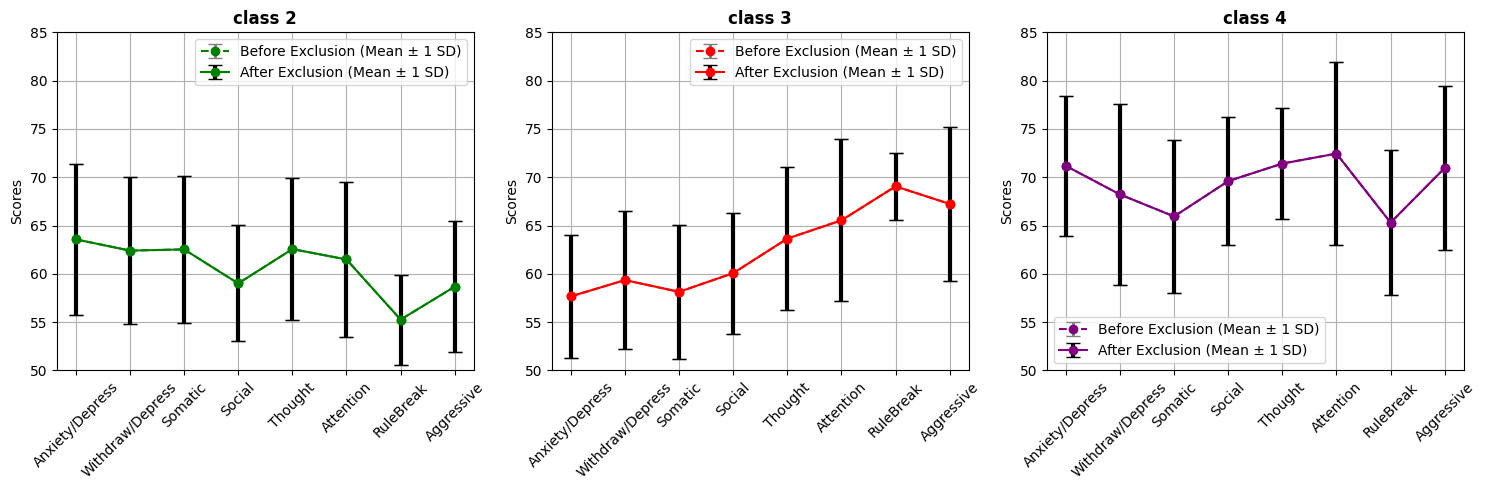

Figure S7. Comparisons in proportions of subjects with clinically significant syndrome across the CBCL syndrome scales before and after excluding high entropy subjects ($E>0.2$).

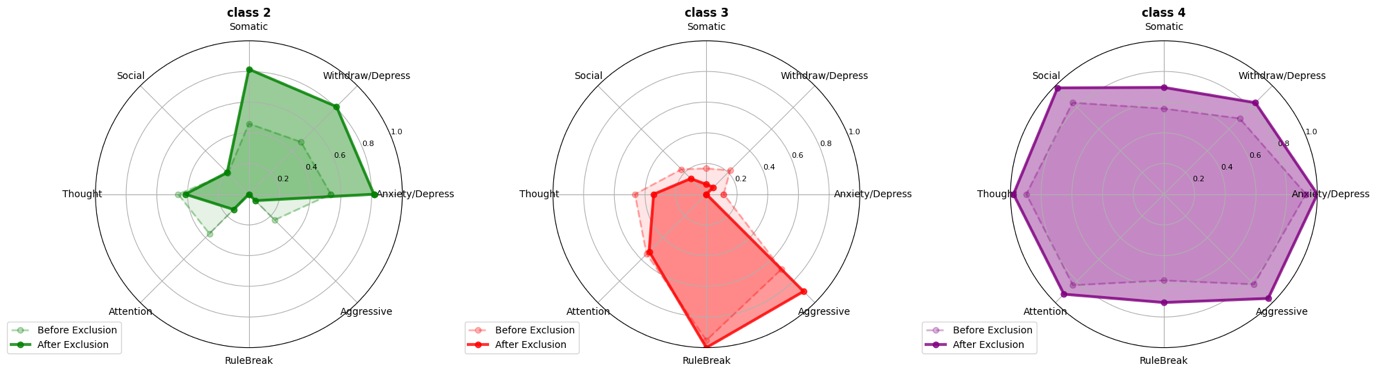

Figure S8. Change in mean and variation of syndrome scores across the CBCL syndrome scales of class 2-4 before and after excluding high entropy individuals ($E>0.2$).

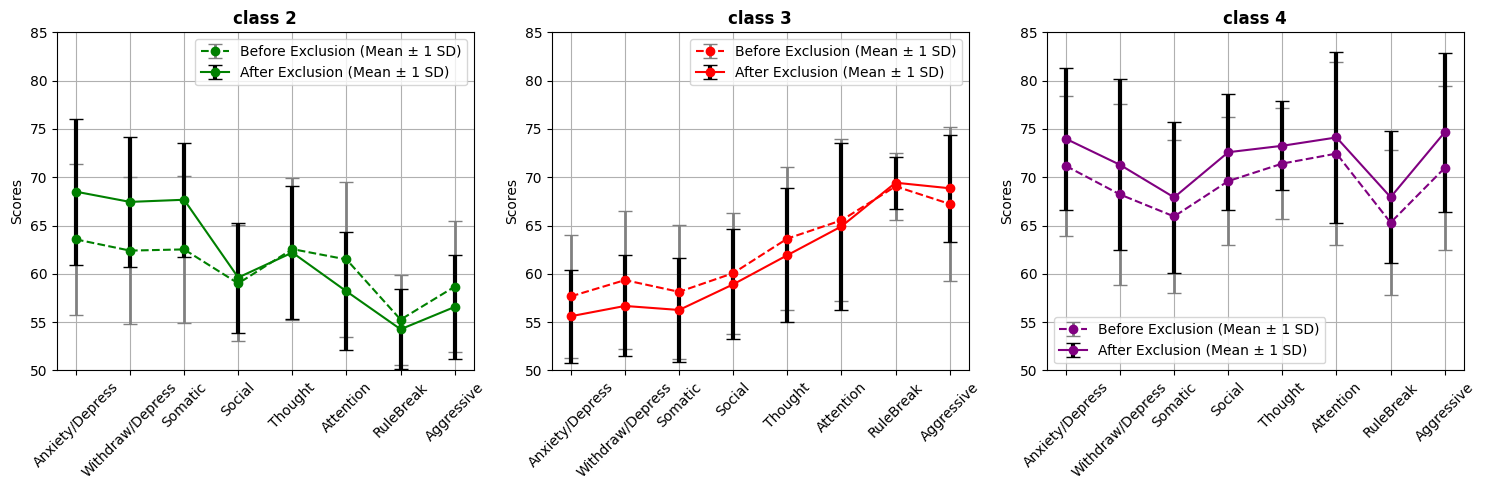

Figure S9. Subject-level syndrome patterns across the CBCL syndrome scales of psychopathological cohorts.

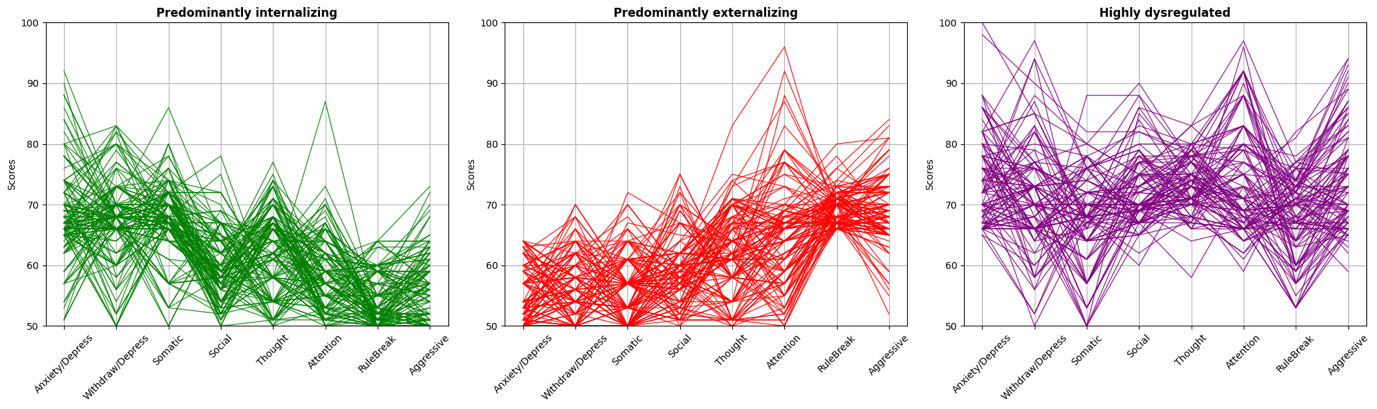

Table S6. Demographics of the psychopathological cohorts

|  | **Predominantly internalizing** | **Predominantly externalizing** | **Highly dysregulated** |
| --- | --- | --- | --- |
| **N** | 135 | 125 | 102 |
| **Biological sex, n (%)** |  |  |  |
| Male | 65 (48.1%) | 73 (58.4%) | 66 (64.7%) |
| Female | 70 (51.9%) | 52 (41.6%) | 36 (35.3%) |
| **Age in months** |  |  |  |
| mean (SD) | 119.83 (6.94) | 119.28 (7.40) | 119.29 (7.33) |
| **Race/ethnicity, n (%)** |  |  |  |
| White | 68 (50.4%) | 49 (39.2%) | 42 (41.2%) |
| Black | 9 (6.7%) | 43 (34.4%) | 20 (19.6%) |
| Hispanic | 39 (28.9%) | 10 (8.0%) | 23 (22.5%) |
| Asian | 1 (0.7%) | 0 (0.0%) | 0 (0.0%) |
| Other | 18 (13.3%) | 23 (18.4%) | 16 (15.7%) |
| **Annual income in $, n (%)** |  |  |  |
| < 5,000 | 1 (0.7%) | 12 (9.6%) | 14 (13.7%) |
| 5,000 - 11,999 | 4 (3.0%) | 15 (12.0%) | 9 (8.8%) |
| 12,000 - 15,999 | 9 (6.7%) | 5 (4.0%) | 5 (4.9%) |
| 16,000 - 24,999 | 5 (3.7%) | 8 (6.4%) | 10 (9.8%) |
| 25,000 - 34,999 | 16 (11.9%) | 17 (13.6%) | 11 (10.8%) |
| 35,000 - 49,999 | 20 (14.8%) | 7 (5.6%) | 9 (8.8%) |
| 50,000 - 74,999 | 19 (14.1%) | 14 (11.2%) | 12 (11.8%) |
| 75,000 - 99,999 | 13 (9.6%) | 5 (4.0%) | 6 (5.9%) |
| 100,000 - 199,999 | 29 (21.5%) | 24 (19.2%) | 11 (10.8%) |
| > 200,000 | 9 (6.7%) | 5 (4.0%) | 3 (2.9%) |
| Income Not Provided | 10 (7.4%) | 13 (10.4%) | 12 (11.8%) |

Table S7. Shapiro-Wilk test results for the deviation distribution of each cohort for each cortical modality (Shapiro-Wilk W-statistic (*p*)). HCs refers to healthy controls, PI refers to the Predominantly internalizing cohort, PE refers to the Predominantly externalizing cohort, HD refers to the Highly dysregulated cohort.

| Cohort | Cortical Thickness | Cortical Volume | Cortical Surface Area |
| --- | --- | --- | --- |
| HCs | 0.86 (p = 7.40e-17) | 0.74 (p = 1.23e-22) | 0.87 (p = 2.84e-16) |
| PI | 0.83 (p = 3.23e-10) | 0.92 (p = 2.65e-06) | 0.92 (p = 5.15e-06) |
| PE | 0.80 (p = 4.93e-10) | 0.78 (p = 1.37e-10) | 0.91 (p = 5.57e-06) |
| HD | 0.85 (p = 1.55e-07) | 0.78 (p = 1.47e-09) | 0.92 (p = 6.54e-05) |

Figure S10. Bootstrapped 99% confidence intervals of effect size (Cliff’s delta) for cortical region deviation scores in cortical thickness, comparing psychopathological cohorts to Healthy Controls. Only features with significant positive deviations are shown.

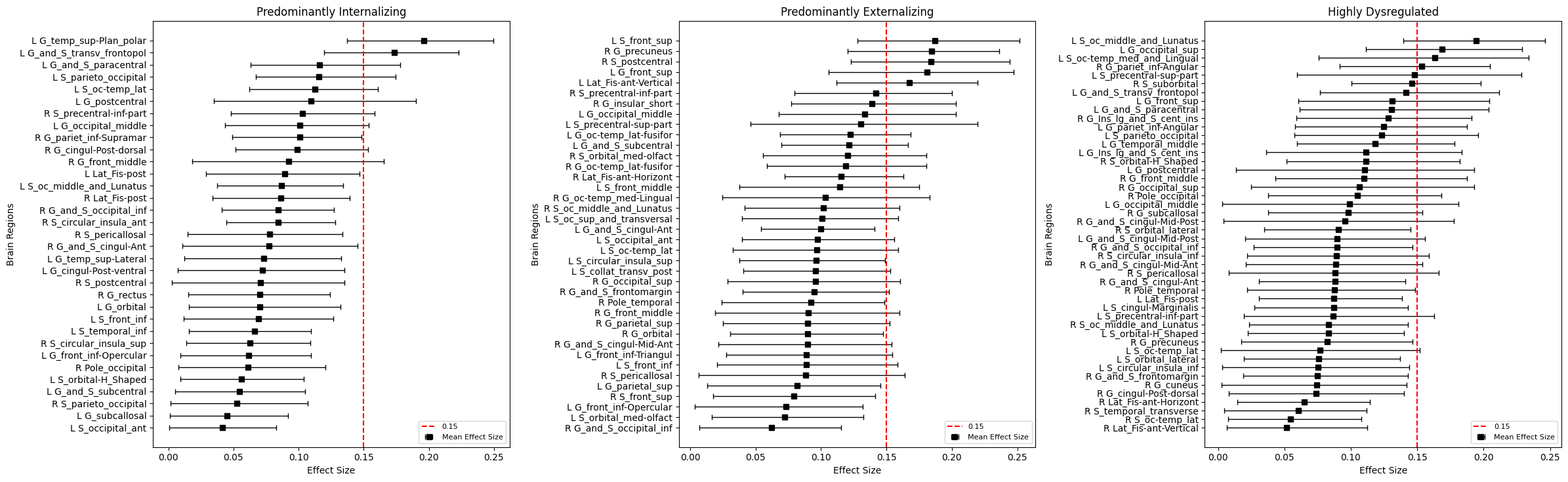

Figure S11. Bootstrapped 99% confidence intervals of effect size (Cliff’s delta) for cortical region deviation scores in cortical volume, comparing psychopathological cohorts to Healthy Controls. Only features with significant positive deviations are shown.

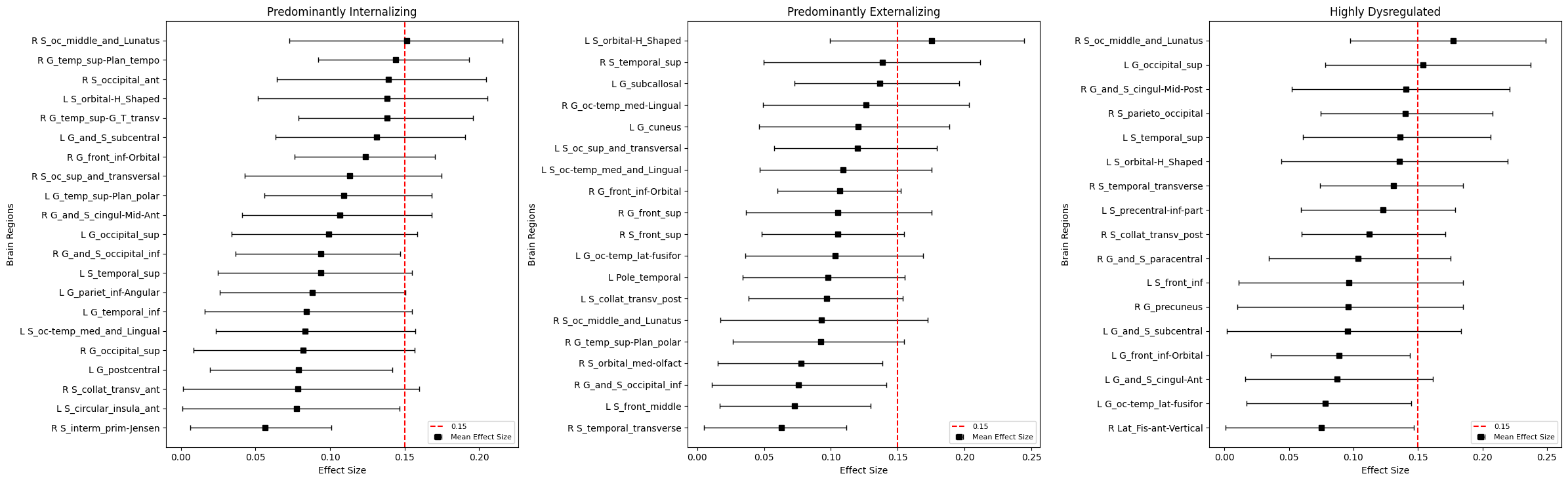

Figure S12. Bootstrapped 99% confidence intervals of effect size (Cliff’s delta) for cortical region deviation scores in cortical surface area, comparing psychopathological cohorts to Healthy Controls. Only features with significant positive deviations are shown.

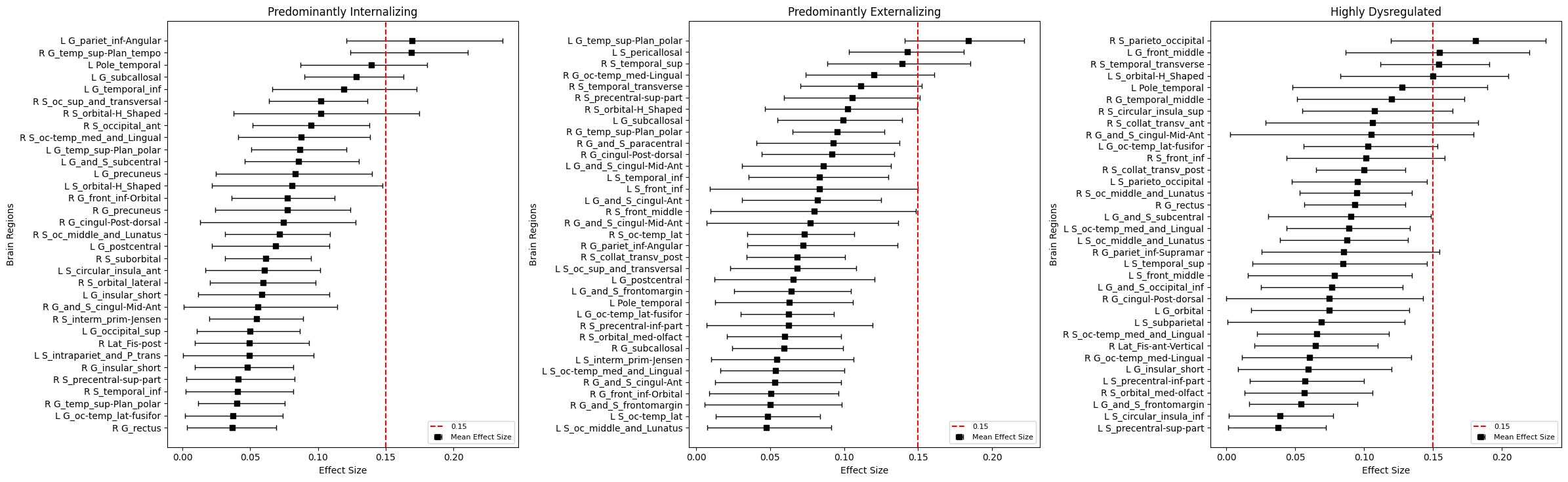

Figure 13. Comparisons in whole cortex deviation distributions of the latent classes 2-4 against HCs in terms of cortical thickness, volume and surface area. Within each modality Mann-Whitney U tests with Benjamini-Hochberg FDR correction (correcting for three comparison tests within each modality) assessed statistical significance: * for $FDR corrected at p < 0.05$, ** for $FDR corrected at P< 0.01$, *** for $FDR corrected at p < 0.001$.

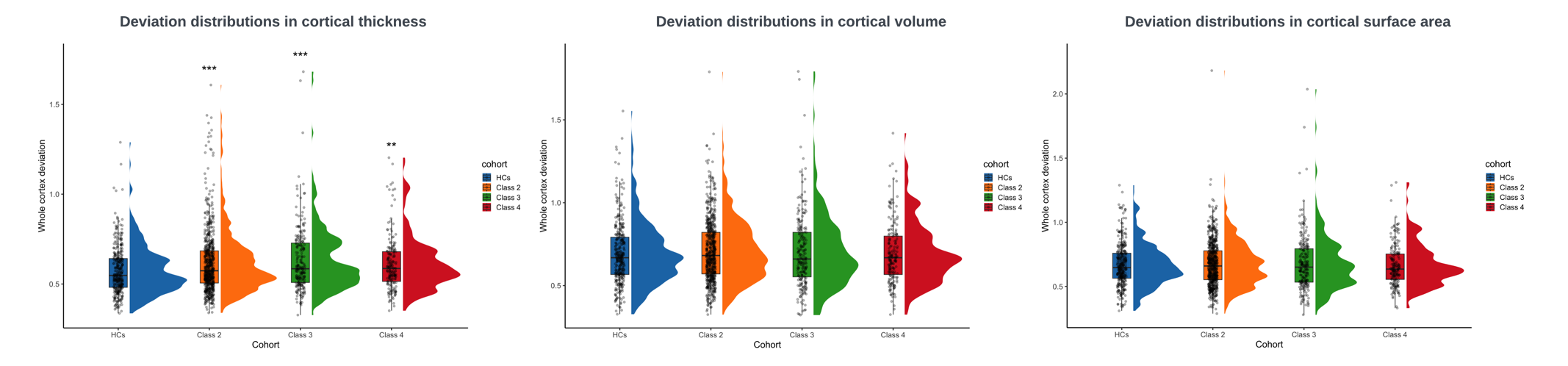

Figure S14. Bootstrapped 99% confidence intervals of effect size (Cliff’s delta) for cortical region deviation scores in cortical thickness, comparing original latent classes to Healthy Controls. Only features with significant positive deviations are shown.

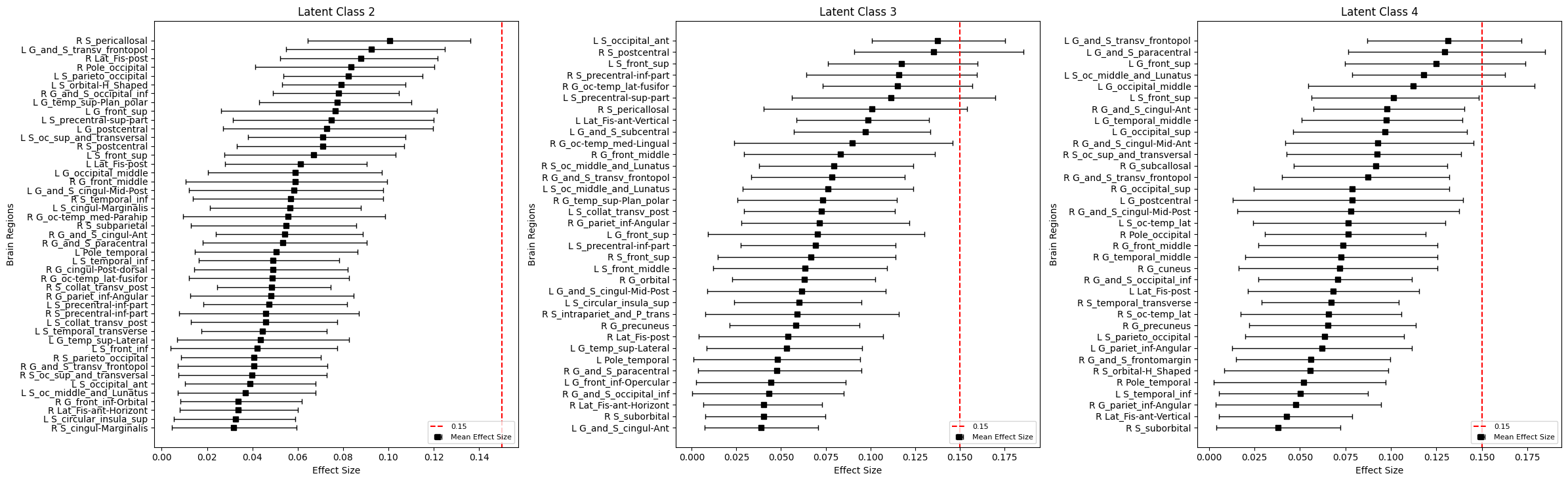

Figure S15. Bootstrapped 99% confidence intervals of effect size (Cliff’s delta) for cortical region deviation scores in cortical volume, comparing original latent classes to Healthy Controls. Only features with significant positive deviations are shown.

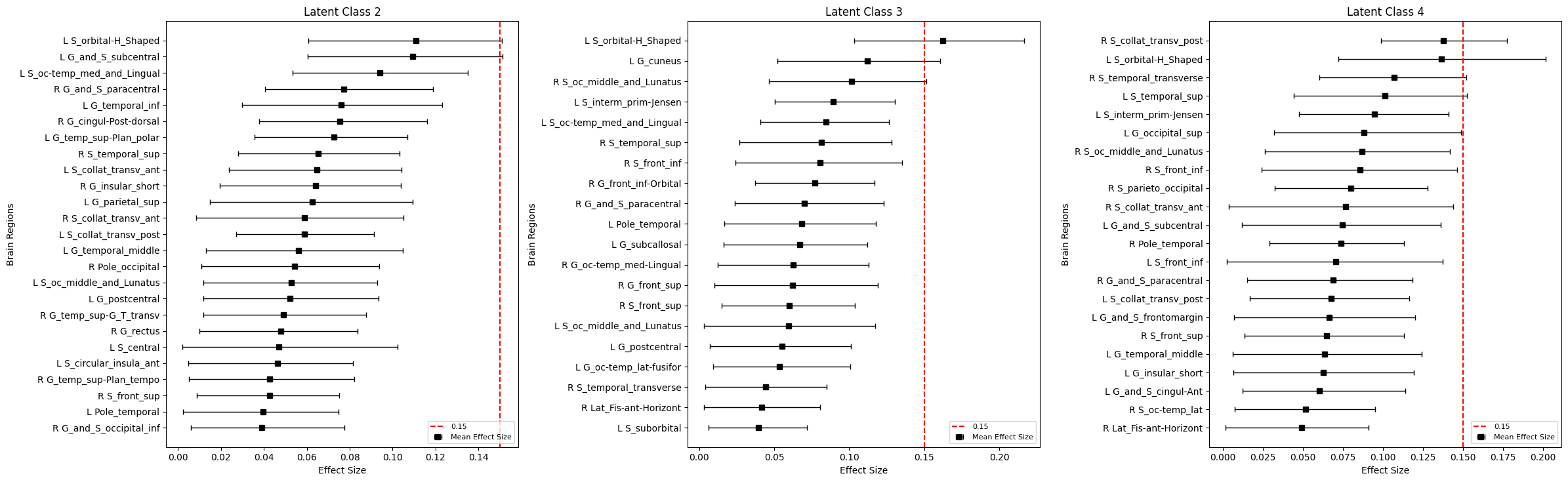

Figure S16. Bootstrapped 99% confidence intervals of effect size (Cliff’s delta) for cortical region deviation scores in cortical surface area, comparing psychopathological cohorts to Healthy Controls. Only features with significant positive deviations are shown.

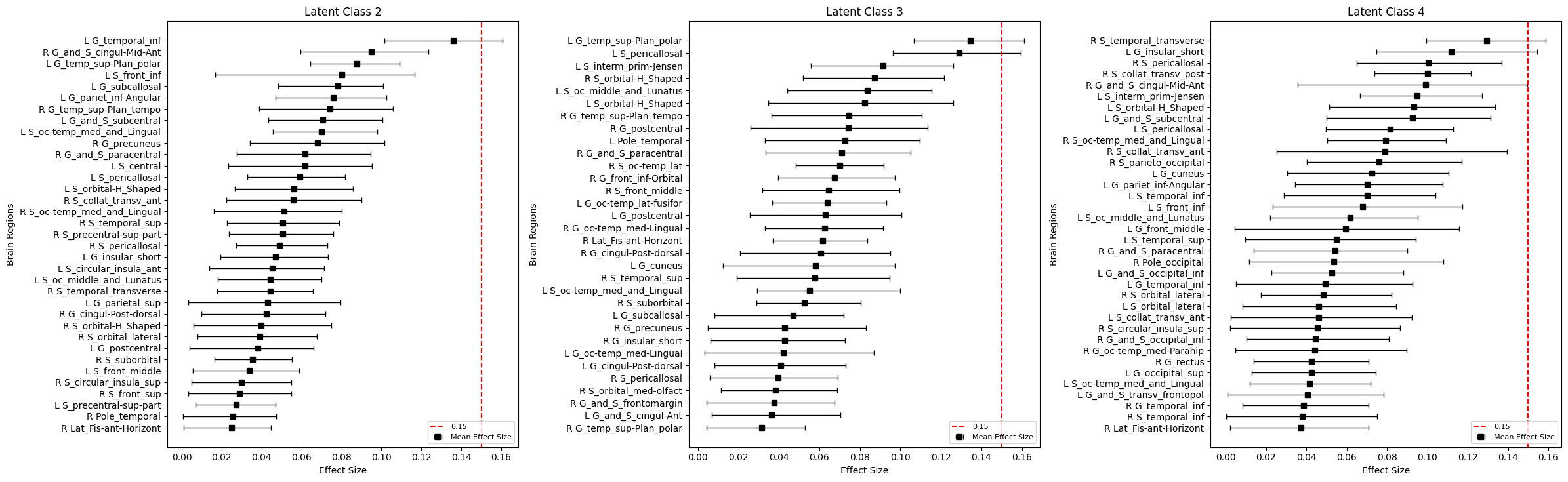
